# Decreased intra-lymphocyte cytokines measurement in septic shock patients: a whole blood proof of concept study

**DOI:** 10.1101/158089

**Authors:** Wilfried Letessier, Julie Demaret, Morgane Gossez, Camille Allam, Fabienne Venet, Thomas Rimmelé, Guillaume Monneret

## Abstract

Functional testing protocols are thought to be the gold standard for the exploration of the immune system. However, in terms of routine analysis, they present numerous drawbacks and consequently their use is mainly limited to research applications. In the clinical context of septic shock, characterized by marked lymphocyte alterations, a new approach for lymphocyte intracellular cytokine measurement in whole blood upon was evaluated in a proof-of-concept study. Following lymphocyte activation, simultaneous intracellular labeling of Interferon-γ (IFN-γ), Tumor Necrosis Factor-α (TNF-α), and Interleukin-2 (IL-2) was performed in CD4^+^ and CD8^+^ T cells (identified by surface marking). The analysis was carried out by flow cytometry (6 colors). Results obtained in septic patients (n = 22) were compared to those of healthy volunteers (n = 8). Independently of lymphopenia, there were significant differences between groups. In particular there was significant decrease in the production of IL-2 and TNF-α in septic patients, while the production of IFN-γ was not significantly altered. Polyfunctional results showed that patients presented with increased percentages of triple negative lymphocytes. In contrast, volunteers had higher proportions of triple positive cells. The approach could be performed in a robust and consistent way, taking 4.5 hours to complete. Moreover, clear differences could be observed between clinical groups with this modified method. These characteristics illustrate the potential of this novel whole blood protocol for clinical applications. However, further research is required to determine the applicability compared to alternative test and to evaluate clinical performances in larger cohorts of patients.

## 1. INTRODUCTION

In clinical immunology laboratories, functional testing remains the gold standard because it directly measures the capacity of a cell population to respond to an immune challenge (1,2). However, although providing excellent insights regarding pathophysiology, this approach remains barely usable for routine clinical monitoring due to several drawbacks (*e.g.*, long incubation time, lengthy cell purification procedures, cell permeabilization, several staining steps, and numerous wash cycles). As a consequence, these protocols remain difficult to be standardized and thus they do not easily fulfil criteria for certification and accreditation of clinical laboratories (3). Thus, a major challenge is therefore to bring functional testing to routine clinical practice.

Among immune functional assays, measurement of intracellular cytokines by flow cytometry appears as one of the most helpful (4). It is able to determine both cell functionality and phenotype, while enumerating them. For instance, it is frequently used to characterize immune responses to infectious diseases, assess pyrogenicity of solutions, and measure immunogenicity during clinical trials of vaccines (4-6). However, this assay remains time-consuming (about 6-8 hours for completion, including several technical steps) and is consequently not used in clinical monitoring except for rare conditions (*e.g.*, immune deficiencies, (1,7)).

We recently reported a novel approach to intracellular staining for flow cytometry that uses whole blood, and which is both rapid and robust (7). Based on a similar approach, the aim herein was to assess simultaneous measurement of TNF-α, IFN-γ, and IL-2 in CD4 and CD8 T cells in clinical samples. This proof of concept study was performed in the clinical context of septic shock in which T cells are believed to have an exhausted phenotype characterized by decreased cytokine production capacity (8).

## 2. MATERIALS AND METHODS

### 2.1 Study population

Twenty-two septic shock patients (according to the diagnostic criteria of the International Guidelines for Management of Severe Sepsis and Septic Shock (9)) admitted to the surgical ICU of E. Herriot Hospital (Lyon University hospitals) were enrolled in the study. This work is part of a comprehensive study of immune dysfunctions induced in ICUs which has been submitted to an ethics committee and approved by the institutional review board and registered with the French Ministry of Higher Education and Research (# DC-2008-509). Blood was collected in heparin coated tubes. Clinical and biological data were collected during the follow-up period (until 28 days)

### 2.2 Research reagents

Tubes (12×75 mm) containing a dry coating of a mixture of Phorbol 12-Myristate 13-Acetate (PMA) and Ionomycin to stimulate the production of cytokines by T cells, and Brefeldin A (BrefA) to block cytokine secretion via the Golgi apparatus. In addition, tubes (12×75 mm) containing a dry coating of conjugated antibodies formulated for this assay were also used: Fluorescein Isothiocyanate (FITC)- labeled anti-IFNγ (clone 45.15), Phycoerythrin (PE)-labeled anti-TNF-α (clone IPM2), PE-Cyanine 7 (PC7(-labeled anti-IL-2 (clone MQ1-17H12), Alexa Fluor 700 (A700)-labeled anti-CD8 (clone B9.ll), Alexa Fluor 750 (A750)-labeled anti-CD3 (UCHT1) and Pacific Blue (PB)-labeled anti-CD4 (13B8.2). These reagents are custom-made, optimized for this study by Beckman Coulter Immunotech (Marseille, France).

### 2.3 Intracellular staining procedure

A schematic protocol representation is presented in Figure 1. Briefly, 50 microliters of undiluted whole blood was directly added to the stimulation tube (PMA-Ionomycin-BrefA), or to an empty control tube, and incubated 3 hours at 37°C. Samples were then treated according to the regular PERFIX-no centrifuge (nc) procedure (Beckman Coulter, Brea, CA, US). Briefly, 25 µL of PERFIX-nc fixative reagent was added and the sample incubated for 15 minutes at room temperature in the dark, 2 mL PBS was then added and the tubes centrifuged at 150g for 5 minutes at 10°C, the supernatant was aspirated, and 25 µL of FBS were added to the pellet. Samples were then concomitantly permeabilized and stained for 45 minutes at room temperature in the dark (dried antibodies were resuspended extemporaneously with 300 µL permeabilizing reagent).The cells were then washed with 3 mL PERFIX-nc final reagent. The pellet was finally resuspended by adding 500 µL of PERFIX-nc final reagent. The entire protocol needed approximately 4.5 h to be completed.

**Figure 1:**
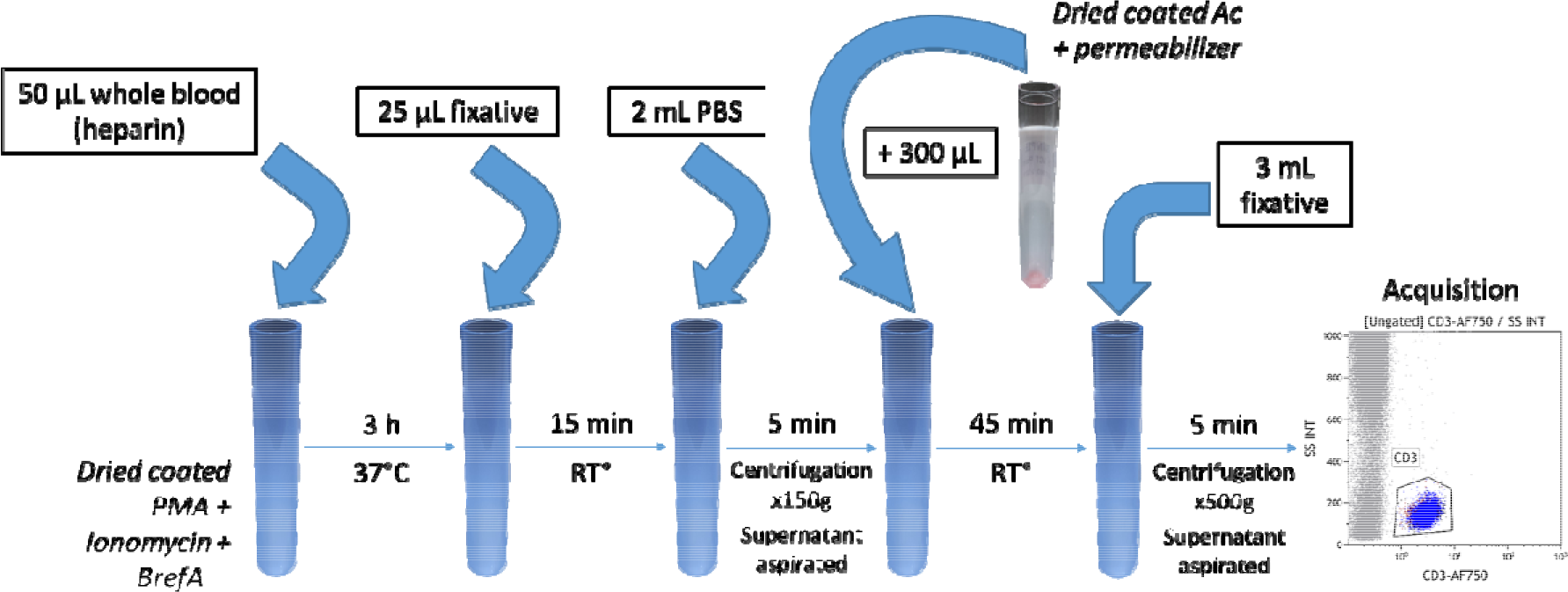
Schematic representation of the staining procedure.

### 2.4 Flow cytometry data acquisition and analysis

Data acquisition was performed on a Navios Flow Cytometer (Beckman Coulter). Our instrument was daily calibrated with Flow-check Pro Fluorospheres (Beckman Coulter) to control optical alignment and fluidic system and Sphero Rainbow Calibration (8 peaks) beads (Spherotech, IL, USA) to ensure MFI stability overtime. T cells were first gated out from other cells on the basis of labelling with CD3. Within the CD3^+^ T cell population, CD4^+^ T cells and CD8^+^ T cells were identified based on CD4/SSC and CD8/SSC dot-plots. Illustrative gating strategy is shown in Figure 2. Intracellular TNF-α, IFN-γ, and IL-2 expressions were then measured on CD4^+^ and CD8^+^ T cell subpopulations. All results were expressed either as percentages of cytokine-positive T cells (% positive cells) among the total CD4^+^ or CD8^+^ T cell subpopulations (positivity threshold was defined based on non-stimulated values from healthy donors and set up at first decade), or as MFI of the entire T cell subpopulation (Figure 2C). Polyfunctional analysis was performed with tree function of Kaluza software (Beckman Coulter).

**Fig 2:**
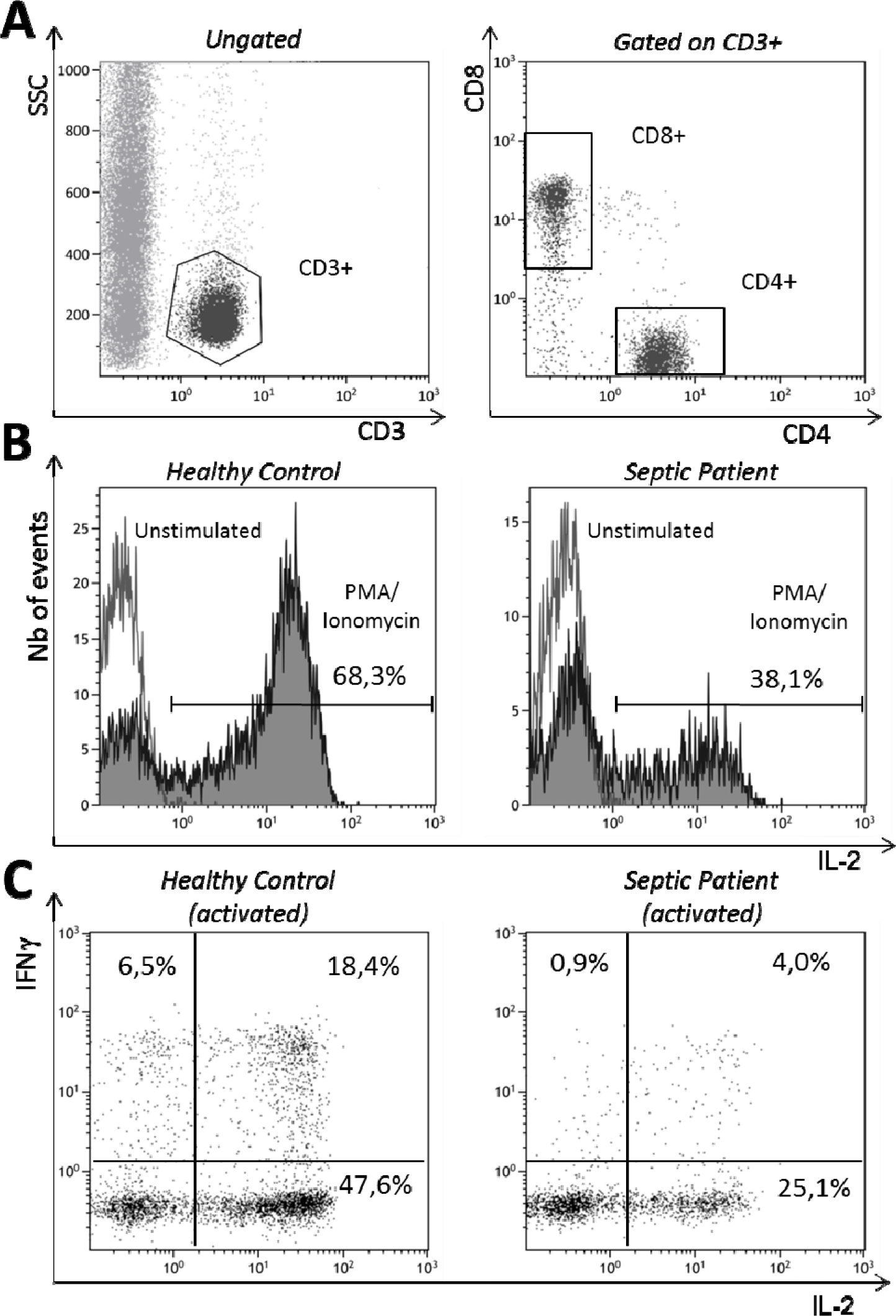
Gating stategy and illustrative examples of intracellular cytokine production measurement in T-cells subsets. A) CD3+ lymphocytes were identified based on side-scattered (SSC) versus CD3 expression on a biparametric histogram (left). CD4+ and CD8+ were identified among selected CD3+ cells (right). (B) Overlay representation of CD4+ intracellular IL-2 measurement in non-stimulated conditions (white) or after a 3 hour-stimulation with phorbol 12-myristate 13-acetate (PMA) and lonomycin (grey) in a Healthy Control (left) or a Septic Patient (right). (C) Biparametric histograms, representing IL-2 and IFNy production, in CD4+ after stimulation, in a Healthy Control (left) or a Septic Patient (right).

### 2.5 Statistical analysis

Results (% positive cells and MFI) were expressed as individual values and medians +/- IQR (interquartile range). Results for the three cytokines (TNF-α, IFN-γ, and IL-2) in both T cell subpopulations (CD4^+^ and CD8^+^) with or without PMA-Ionomycin challenge among healthy volunteers and septic shock patients were compared, using the non-parametric Mann-Whitney U test or Wilcoxon paired test. Correlations were investigated using the Spearman correlation test. Statistical analyses were performed with GraphPad Prism^®^ software (version 5.0; GraphPad Software, La Jolla, CA, US). A p-value < 0.05 was considered as statistically significant.

## 3. RESULTS

### 3.1 Patients and healthy volunteers

Twenty-two septic shock patients (sampled during first 48 hours after diagnosis) and eight healthy controls were included. Five patients were sampled a second time at D3-4. At onset of shock, the median [Interquartile range, IQR] age was 74 years [68-83], Simplified Acute Physiology Score (SAPS) II was 59 [IQR 47.5-74], and the Sequential Organ Failure Assessment score was 8 [IQR 7-11.5]. Seven patients (32%) died within 28 days and secondary nosocomial infections were diagnosed in 4 patients (18%). At D3-4, septic shock patients presented with usual characteristics of injury-induced immunosuppression with a reduced mHLA-DR expression and low CD4^+^ lymphocytes count in comparison with normal values (Table 1). Healthy volunteers (4 women / 4 men, from 48 to 60 years old) all presented with normal values regarding mHLA-DR (median: 18 500 ABC) or CD4^+^ lymphocyte count (median: 798 cells/µL).

**Table 1:**
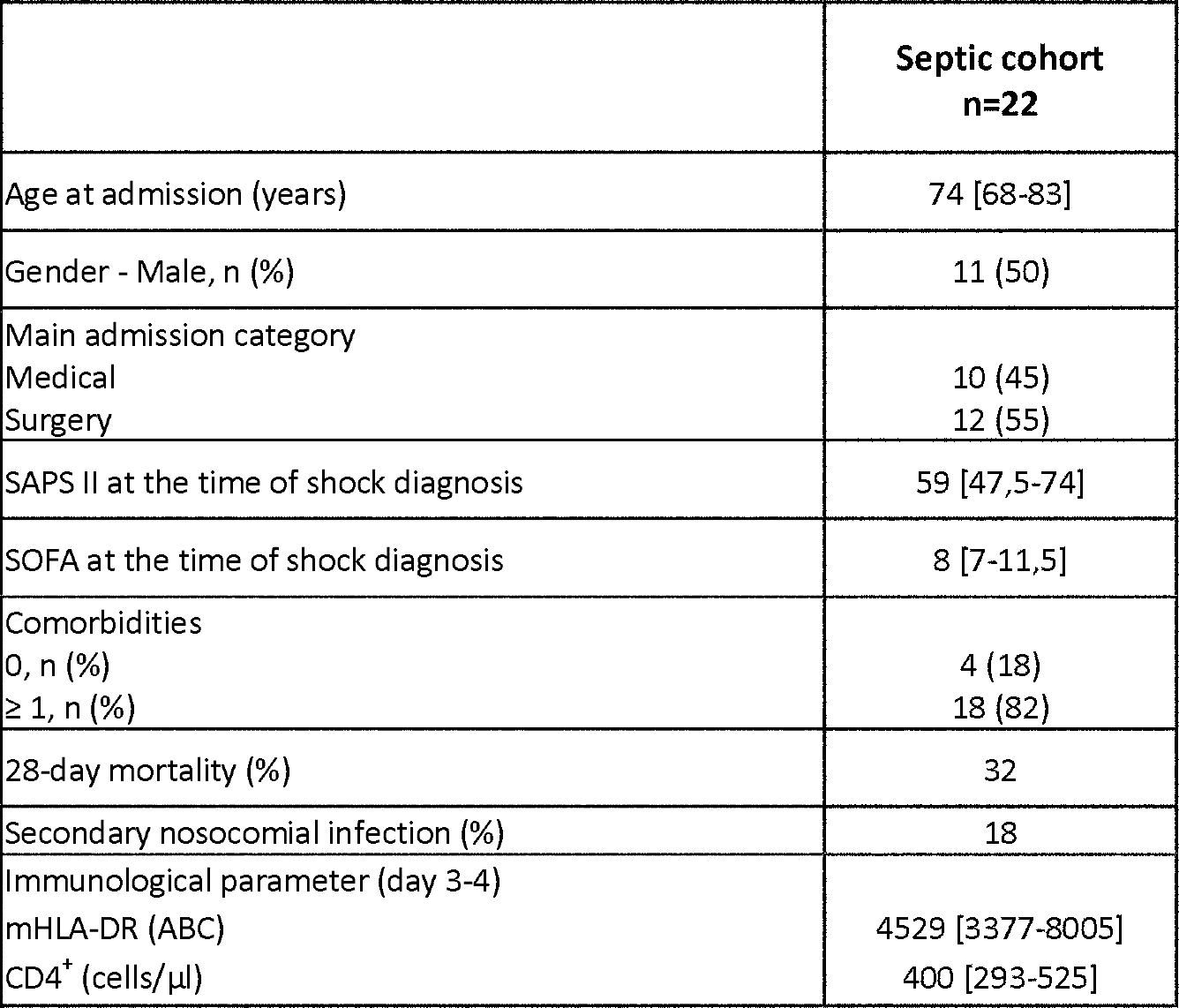
Clinical and immunological data for patients with septic shock. Numeric variables are presented as median and IQR (quartile 1 to quartile 3). Categorical variables are presented as number of cases, and percentages among the total of patients in parentheses. SAPS II: Simplified Acute Physiology Score II, calculated at the patient inclusion. SOFA: Sequential Organ Failure Assessment, measured after 24h of ICU stay. mFILA-DR: monocyte FILA-DR expression (ABC: antibodies bound per cell, normal values > 15 000), CD4+: absolute value of CD4 T cells (normal values: 500-1250) - measured at days 3-4 after septic shock onset.

|  | Septic cohort<br>n=22 |
| --- | --- |
| Age at admission (years) | 74 [68-83] |
| Gender - Male, n (%) | 11 (50) |
| Main admission category |  |
| Medical | 10 (45) |
| Surgery | 12 (55) |
| SAPS II at the time of shock diagnosis | 59 [47,5-74] |
| SOFA at the time of shock diagnosis | 8 [7-11,5] |
| Comorbidities |  |
| 0, n (%) | 4 (18) |
| ≥ 1, n (%) | 18 (82) |
| 28-day mortality (%) | 32 |
| Secondary nosocomial infection (%) | 18 |
| Immunological parameter (day 3-4) |  |
| mHLA-DR (ABC) | 4529 [3377-8005] |
| CD4<sup>+</sup> (cells/μl) | 400 [293-525] |

### 3.2 Intracellular cytokines

At D1-2 expression of TNF-α, IFN-γ, and IL-2 in both T cell subpopulations was statistically higher following PMA-Ionomycin challenge compared to non-stimulated control, either in healthy subjects or patients (p<0.0001). This was observed when results were expressed either as a proportion of positive cells (Figure 3) or MFI (Figure 4). The proportion of CD4 T cells expressing TNF-α (p = 0.0002) and IL-2 (p = 0.0007) as well as the proportion of CD8^+^ T cells expressing IL-2 (p = 0.0011) after stimulation were significantly lower among septic shock patients than healthy volunteers (Figure 3). Similarly, significant differences were found following PMA-Ionomycin stimulation between patients and healthy volunteers for TNF-α and IL-2 MFI of both CD4^+^ (p = 0.0106, and p = 0.0121, respectively) and CD8^+^ T cells, (p = 0.0157 and p = 0.0053, respectively; Table 4). No significant difference was observed in the kinetic study between patients sampled both at D1-2 and D3-4 (data not shown). Polyfunctional results (from tree analysis, Figure 5A) are exhaustively presented in Table 2. Illustrative examples are provided in Figure 5B. For instance, septic patients presented with increased percentages of triple negative CD4^+^ T cells (p: 0.0001). In contrast, volunteers had more cells simultaneously positive for the 3 measured cytokines both in CD4^+^ T cells (p: 0.009) and CD8^+^ T cells (p: 0.003). We did not find any association between polyfunctional results and mortality.

**Figure 3:**
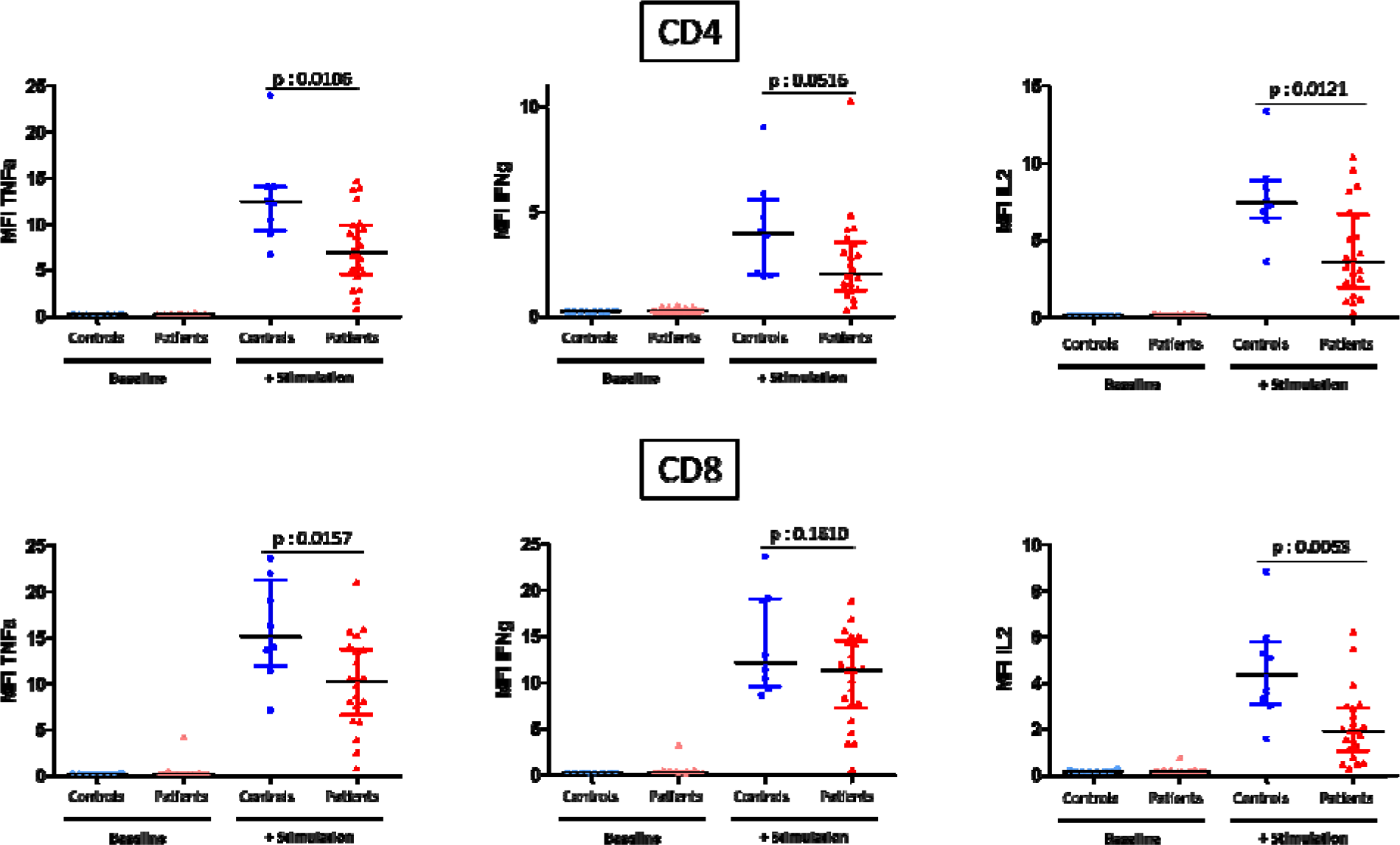
CD4+ and CD8+ T lymphocytes expressing TNF-α, IFN-γ, and IL-2 (as  *%*  of positive cells) in healthy volunteers (n = 8) and septic shock patients (n = 22, days 1-2). Results are represented as individualized points along with the median and IQR.

**Figure 4:**
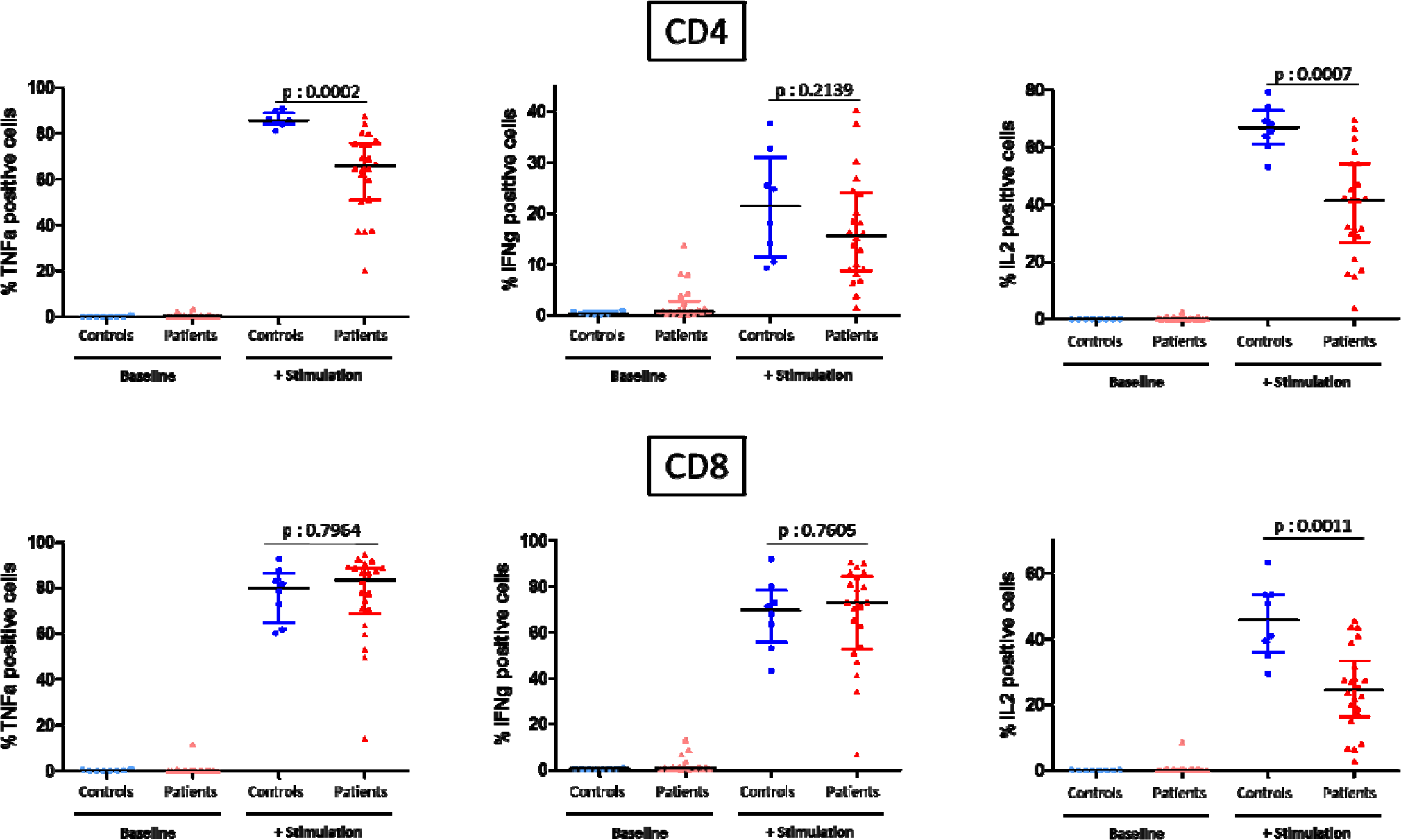
CD4+ and CD8+ T lymphocytes expressing TNF-α, IFN-γ and IL-2 (as mean of fluorescence intensities) in healthy volunteers (n = 8) and septic shock patients (n = 22, days 1-2). Results are represented as individualized points along with the median and IQR.

**Table 2:** Polyfunctional analysis of 3 cytokines intracellular staining. Percentages of cells (among CD8+ and CD4+ populations) producing none, one, two or three cytokines (after PMA / lonomycin stimulation) in controls (n=8) and septic patients (n=22) are presented. Results are expressed as percentages of positive cells among gated cells (i.e., CD4 or CD8 cells) and as medians +/- IQR.

| Cytokine |  | CD4+ |  | CD8+ |  |
| --- | --- | --- | --- | --- | --- |
| Positive : + | Negative : - | Healthy volunteers | Septic patients | Healthy volunteers | Septic Patients |
| IFNγ | - | | | | |
| IL2 | - | 5.5 % [4.8-5.7] | 20.2 % [11.8-35.4] | 13.4 % [9.93-22.13] | 14.4 % [9.0-27.6] |
| TNFα | - | | | | |
| IFNγ | - | | | | |
| IL2 | - | 20.5 % [15.7-23.4] | 24.9 % [21.1-30.1] | 4.7 % [2.6-7.3] | 3.9 % [1.9-9.7] |
| TNFα | + | | | | |
| IFNγ | - | | | | |
| IL2 | + | 2.0 % [1.6-2.3] | 2.3 % [1.9-3.3] | 2.9 % [2.0-3.9] | 1.0 % [0.3-1.7] |
| TNFα | - | | | | |
| IFNγ | + | | | | |
| IL2 | - | 0.1 % [0-0.2] | 0.2 % [0.1-0.4] | 0.7 % [0.6-1.2] | 1.1 % [0.9-2.8] |
| TNFα | - | | | | |
| IFNγ | - | | | | |
| IL2 | + | 47.9 % [42.6-54.7] | 31.6 % [18.4-43.0] | 6.5 % [2.9-9.8] | 4.8 % [1.9-8.4] |
| TNFα | + | | | | |
| IFNγ | + | | | | |
| IL2 | - | 3.0 % [1.4-6.5] | 4.1 % [1.9-7.0] | 29.4 % [16.8-39.1] | 43.5 % [35.0-53.3] |
| TNFα | + | | | | |
| IFNγ | + | | | | |
| IL2 | + | 0.0 % [0-0] | 0.1 % [0-0.2] | 1.0 % [0.5-1.4] | 0.3 % [0.1-0.7] |
| TNFα | - | | | | |
| IFNγ | + | | | | |
| IL2 | + | 17.6 % [10.3-23.0] | 8.4 % [4.9-13.9] | 37.9 % [31.6-43.7] | 20.2 % [11.2-33.2] |
| TNFα | + | | | | |

**Figure5:**
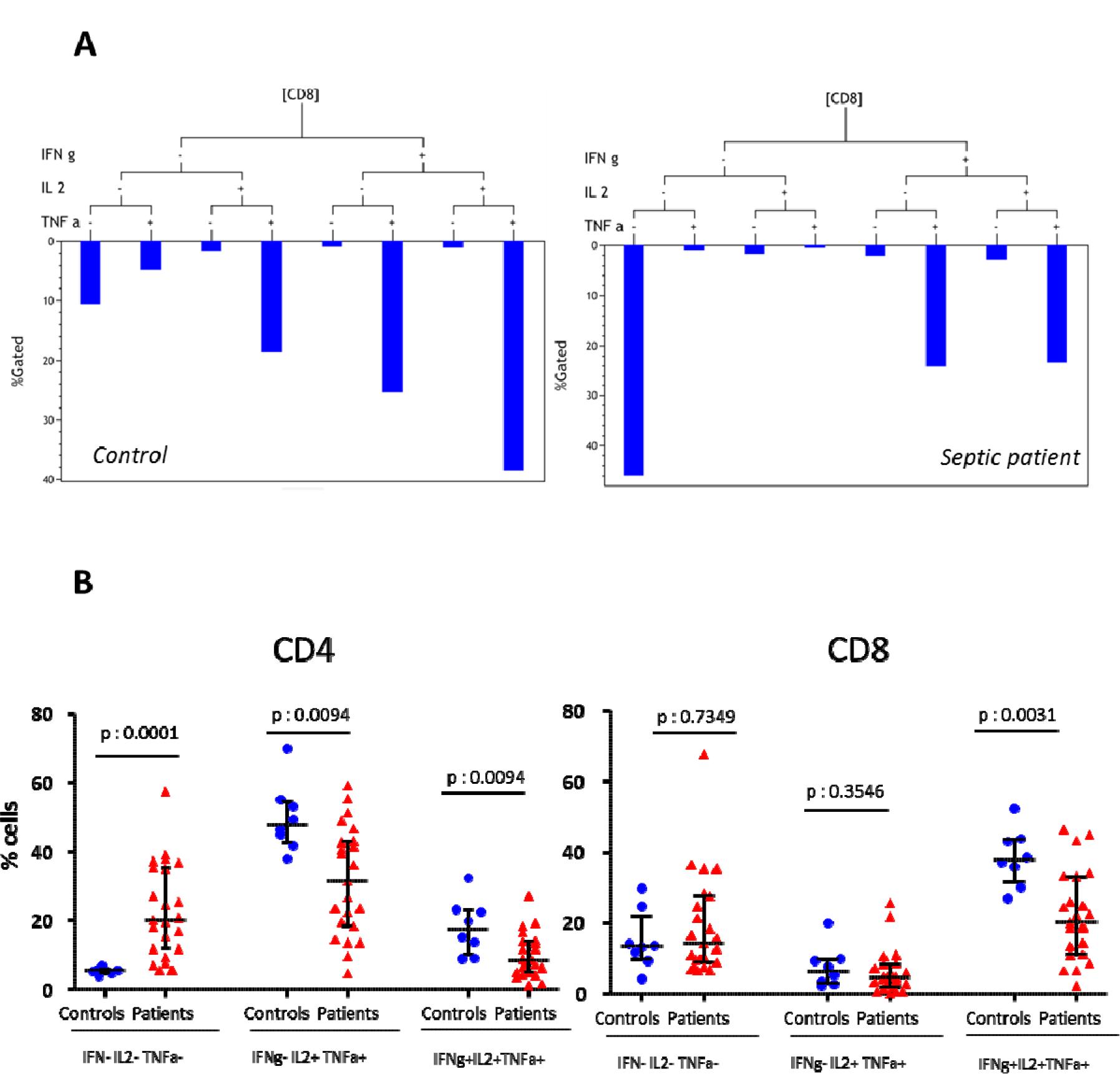
Polyfunctional analysis of 3 cytokines intracellular staining. (A) Representative examples of tree analysis in CD8+ cells from a healthy volunteer and a septic patient. Each bar represents the percentage of cells (among CD8+ population) producing none, one, two or three cytokines. (B) Polyfunctional cytokine expression between controls (n=8) and septic patients (n=22) in CD4+ T-cells (left) and CD8+ T-cells (right). Illustrative results are represented for 3 possibilities (IFN-/IL2-/TNF-), (IFN-/IL2+/TNF+), (IFN+/IL2+/TNF+) as individualized points along with the median and IQR (p values for comparison according to Mann Whitney test).

### 3.3 Correlations

We did not find any significant correlations between CD4 and CD8 T cell counts and intracellular cytokine expressions (% positive cells and MFI). There was only one significant correlation (IL-2 production and CD8 T cell count) and it was negative (r: −0.46, p: 0.034). These results mostly highlight no major significant correlation between lymphopenia and lymphocyte function. No correlation was found with decreased MFILA-DR (data not shown) or SAPS II. Slightly significant correlations were observed between SOFA score and TNF (r: −0.53, p: 0.01) or IL-2 (r: −0.52, p: 0.01) when expressed as MFI in CD4 T cells. From polyfunctional results, we also found negative correlations between SOFA score and triple negative CD4 cells (r: 0.55, p: 0.01) or triple negative CD8 cells (r: 0.46, p: 0.01).

## 4 DISCUSSION

In the present work a new protocol for the quantification of intracellular cytokines in T lymphocytes was assessed. The approach, based on whole blood (50 µL), is straightforward and allows the simultaneously measurement of three cytokines (TNF-α, IFN-γ, and IL-2) in subsets defined by the addition of two surface markers (CD4 and CD8). The entire protocol takes approximately 4.5 hours to be completed, which seems to be quicker when compared to the usual published intracellular cytokine detection protocols that take up to 8 hours (4,10,11). In addition, the dry coating of reagents may improve standardization by reducing the pipetting steps and permitting storage at room temperature. Similar whole blood protocols reported in the literature (4,10) have certain limitations compared to the one presently reported: for instance the initial incubation requires 6 hours (2 hours to stimulate cells, followed by 4 hour-incubation to block cytokine secretion with brefeldin) instead of only 3 hours herein, a greater number of washing steps, no dried reagents (including brefeldin) and antibodies, and reagents cannot be stored at room temperature. As we did not perform usual protocol at the same time in the present study, strict comparison of technical aspects remains to be done. That said, with only a few manual steps, the present protocol allows the technician to perform this protocol among other routine tasks. The present results aggregate with those obtained in monocytes regarding TNF-α production (7). Taken together, they tend to highlight practical improvements in intracellular cytokine measurement.

In the delayed step of septic shock, exhaustion of T cells characterized by a loss of function (e.g. proliferation) and decrease in cytokine release is commonly described (8,12). With the present protocol, as expected, cytokine results significantly differentiated septic shock patients from healthy volunteers. Both in CD4^+^ and CD8^+^ T cells, TNF-α and IL-2 synthesis was significantly reduced. Based on the chronology of T cell exhaustion *(i.e.*, the first cytokine production to be stopped is IL-2, then TNF-α, and finally IFN-γ), septic patients in our study seem to be in a rather late phase of the process (loss of IL-2 and TNF-α), but are not in the latest stage since production of IFN-γ is maintained (13). These results could be explained by analyses having been performed 1 or 2 days following the onset of septic shock, while the decrease of IFN-γ production along with impairment of other functions *(e.g.*, proliferation) would occur later. Nevertheless, in few patients with additional samples at D3-4, no significant difference could be further detected. That said, polyfunctional analysis revealed that IFN-γ production may still be partially altered since we observed significant differences between controls and septic patients when focusing on triple positive cells both in CD4 T and CD8 T cells. From a therapeutically perspective, knowing that septic lymphocytes did not enter final stage of exhaustion reinforces the potential of rejuvenating them with IL-7 (8,14).

There was no correlation between the absolute value of CD4^+^ or CD8^+^ T cells and their ability to produce cytokines. This illustrates the independence of lymphocyte functionality from lymphopenia and illustrates the potential interest of using functional testing to finely characterize patients. For instance, on the basis of a polyfunctional protocol, Snyder et al. could predict the risk of developing CMV infection in patients who received a lung transplant (15). That said, the clinical interest of the present approach remains to be assessed.

The study does, however, present certain limitations. Firstly, this test is based on antigen-independent stimulation of the lymphocytes. Thus, flow cytometry may not be suitable for antigen stimulation as assessed by other techniques *(e.g.*, ELISPOT). In addition, as a proof-of-concept study, the main objective of the present work was to assess feasibility of this whole blood protocol in clinical samples. Thus, comparison to other methods for assessing intracellular cytokines production was not made, and this remains to be performed. In line with this proof of concept objective, a relatively small number of patients were included and no conclusions as to clinical outcomes could be made (mortality, occurrence of nosocomial infections). It is particularly true for investigating the potential of polyfunctional analysis that would be likely fully revealed in large cohort of patients allowing robust statistics. This work now deserves to be widely assessed and validated in a larger ICU population, and if possible in multicenter studies.

In conclusion, these preliminary results indicate the feasibility of this novel whole blood protocol. Its simplicity, robustness, and rapidity allow it to be integrated into a routine work load or in the monitoring of clinical trials. Beyond septic shock and ICU patients, it appears reasonable to envisage a wider use in other clinical situations where the assessment of intracellular cytokine synthesis might help in monitoring immunosuppressed patients *(e.g.*, transplantation, cancer, immune deficiencies) or treatments modulating immune responses *(e.g.*, biotherapies).

## 5. ACKNOWLEDGMENTS

This work was supported by Hospices Civils de Lyon and Université Claude Bernard Lyon 1. It was also supported by Beckman Coulter through donations of the research reagents used in this evaluation. Beckman-Coulter made no contribution to the study design, or in the collection or interpretation of the data. Similarly, Beckman Coulter had no role in the preparation of the manuscript or the decision to submit it for publication. Beckman-Coulter product names are trademarks or registered trademarks and are used with permission. The authors would like to thank Anne Portier for technical assistance and Philippe Robinson for English editing.

